# Offset mass carrier proteome improves quantification of multiplexed single cell proteomics

**DOI:** 10.1101/2024.11.08.622689

**Authors:** Tommy K. Cheung, Ying Zhu, Christopher M. Rose

## Abstract

Multiplexed single cell proteomics by mass spectrometry (scpMS) approaches currently offer the highest throughput as measured by cells analyzed per day. These methods employ isobaric labels and typically a carrier proteome - a sample added at 20-500x the single cell level that improves peptide sampling and identification. Peptides from the carrier and single cell proteomes exist within the same precursor isotopic cluster and are co-isolated for identification and quantification. This represents a challenge as high levels of carrier proteome limit the sampling of peptide ions from single cell samples and can potentially lead to decreased accuracy of quantitative measurements. Here, we address this limitation by introducing a triggered by offset mass acquisition method for scpMS (toma-scpMS) that utilizes a carrier proteome labeled with non-isobaric tags that have the same chemical composition but different mass as the labels used for quantitative multiplexing. Within toma-scpMS the carrier proteome and single cell proteome are separated at the precursor level, enabling separate isolation, fragmentation, and quantitation of the single cell samples. To enable this workflow we implemented a custom data acquisition scheme within inSeqAPI, an instrument application programming interface program, that performed real-time identification of carrier proteome peptides and subsequent triggering of offset single cell quantification scans. We demonstrate that toma-scpMS is more robust to high-levels of carrier proteome and offers superior quantitative accuracy as compared to traditional multiplexed scpMS approaches when similar carrier proteome levels are employed.

## Introduction

Single cell proteomics by mass spectrometry (scpMS) enables the unbiased analysis of proteomes within heterogeneous cell populations without the requirement of antibody targeted reagents (1). There are two general approaches to scpMS: label-free and isobaric label based quantitation methods (1).

Label-free methods, such as data-dependent acquisition (DDA) (2) or data-independent acquisition (DIA) (3–5), offer simplified sample preparation and the ability to identify proteins uniquely expressed in rare cell types. For these methods each mass spectrometry (MS) experiment analyzes a single cell with the current potential to analyze 40-80 cells per day (3–5). Recently, multiplexed DIA (mDIA or plexDIA) methods have been introduced that improve throughput by 3-5 fold (6, 7), but more difficult sample preparation as well as increased spectral complexity may limit further increases in multiplexing capacity.

Isobaric label based quantification requires a more complex sample preparation procedure, but currently represents the highest-throughput approach that enables deep proteome coverage through the use of a carrier proteome (8, 9). The first application of scpMS for the analysis of cancer cell lines, SCoPE-MS (10, 11), introduced the concept of a carrier proteome to enable the analysis of single cell proteomes. A carrier proteome is a sample that is the same or similar to the cells being analyzed and is added into a multiplexed experiment at 20-500x the level of the single cells (8, 10–13). Precursor and fragment ions from the carrier proteome overlap, improving sampling and identification of peptides as compared to analyses without a carrier proteome. However, the co-isolation of carrier and single cell peptide ions can challenge the sampling of single cell ions and decrease quantitative accuracy when high levels of carrier proteome are utilized (8, 9). This is due to carrier proteome ions limiting the isolation of single cell ions as well as potential space charge effects that limit detection of single cell reporter ions when a high level of carrier proteome reporter ion is present (8, 9). These effects can be somewhat overcome by increasing the number of ions sampled as well as the maximum time allowed for ion injection (9, 12, 13). However, due to co-isolation of carrier and single cell precursors, increasing the sampling of single cell peptide ions requires additional sampling of carrier proteome ions - exacerbating potential space charging effects.

An alternative approach would be to use an offset mass carrier proteome to avoid the co-isolation of peptides from the carrier and single cell proteomes (14). This can be accomplished by labeling carrier proteome peptides with non-isobaric versions (i.e., TMT-zero, TMT-SH, TMTpro-zero and TMTpro-SH) of the quantitative labels used for single cell multiplexing (i.e., TMT and TMTpro). This enables the carrier proteome peptides to be isolated, fragmented, and identified separately from the isolation, fragmentation, and quantification of single cell peptides. Because single cell peptides are isolated separately, more single cell ions can be sampled, leading to an increase of signal-to-noise within single cell reporter ion channels. This improves quantitative accuracy and decreases the negative effects of high levels of carrier proteome reporter ions. A brief exploration of this approach demonstrated that an offset mass carrier proteome increased signal for bulk diluted ‘single cell’ channels, but that higher levels of carrier proteome led to a decrease in quantified proteins (14). Here, the decrease in protein identifications was likely due to the utilization of a ‘targeted mass difference’ acquisition method that only triggered peptide analysis when both the carrier and single cell precursor ions were detected in an MS1 survey scan. Higher carrier levels make it difficult to detect both the carrier and single cell peptides within the same survey scan, leading to decreased peptide sampling and decreased protein identifications. This implementation also identified ‘single cell’ peptides from the same offset scans used for quantification (14). While this ensures that quantified reporter ions are derived from the identified peptide, this approach does not utilize the carrier proteome fragment ions for peptide identification - a key advantage of the carrier proteome within SCoPE-MS.

We sought to address these challenges by employing a novel intelligent data acquisition (IDA) approach that utilized real-time peptide database search to identify peptides from an offset carrier proteome and trigger subsequent single cell focused scans. IDA methods utilizing real-time peptide identification approaches were introduced more than ten years ago (15, 16), but these implementations required custom instrument code (16) or difficult to use instrument interfaces (15). Recently, the introduction of a real-time database search within the native instrument method editor of Orbitrap Eclipse and Ascend instruments, as well as the growth of programs that utilize the instrument application programming interface (iAPI) to create custom IDA methods has made the utilization of real-time peptide identification more prevalent (17–20). These methods are often focused on multiplexed quantification utilizing isobaric labels because a rapid-ion trap scan can be used to limit collection of slow FTMS quantification scans to peptides that will be identified in a post-run search (17, 20, 21). IDA methods have also been used for scpMS, most notably the RETICLE method that utilizes an ion-trap identification (22) scan to trigger an Orbitrap FTMS2 quantification scan, and pSCoPE which utilizes an advanced inclusion list method to prioritize low-abundance peptides and improve quantification across runs (23).

Here, we describe a novel triggered by offset mass scpMS (toma-scpMS) method within inSeqAPI (20) that utilizes a real-time search to identify carrier proteome peptides and trigger offset mass quantitative analysis of multiplexed single cells peptides. The resulting data enables connection of carrier proteome identification and single cell quantification scans enabling the carrier proteome scan to be used for post-run peptide identification, maintaining a key advantage of the carrier proteome. Through analysis of equimolar and mixed-ratio bulk-dilution samples we compared the SCoPE-MS approach to toma-scpMS and demonstrated that toma-scpMS was more robust to the utilization of higher carrier proteome levels as the number of quantified proteins and the quantification accuracy did not decrease as carrier levels increase. Lastly we utilize toma-scpMS to analyze single cells at varying carrier levels and with FTMS2 or SPS-MS3 quantitation to demonstrate the ability to separate cell types and identify cell type specific proteins.

## Experimental Procedures

### Diluted bulk sample preparation

Experiments analyzing HeLa and K562 samples used commercial tryptic HeLa (Thermo Scientific Pierce) and K562 digests (Promega). Peptide solutions were dried and then reconstituted in 200 mM HEPES (pH 8.5), before being divided into nine (SCoPE-MS) or eleven (toma-scpMS) aliquots. Aliquots were then labeled with TMT (200 mM HEPES pH 8.5, 28% ACN, 2:1 ratio of label:peptide, reaction time of 1 h), with one carrier proteome aliquot being labeled with channel TMT126 (SCoPE-MS) or TMT-SH (toma-scpMS). For SCoPE-MS, quantitative channels were 127n, 128n, 128c, 129n, 129c, 130n, 130c, 131n and channel TMT127c was left empty to circumvent the 13C-isotope signal of the carrier proteome (channel TMT126). For toma-scpMS, equimolar quantitative channels were 126, 127n, 127c, 128n, 128c, 129n, 129c, 130n, 130c, 131n and mixed ratio quantitative channels were 126, 127n, 127c, 128n, 128c, 129n, 129c, 130n. For equimolar mixtures all quantitative channels were mixed to achieve 400 pg on column per channel per injection. For mixed ratio samples, the final four quantitative channels were mixed to achieve 800 pg on column per channel per injection. After labeling, all samples were dried, desalted and then dried again before reconstituting in 0.1% formic acid.

To create samples with varying amounts of carrier proteome, each sample was mixed with the appropriate carrier proteome, such that the carrier proteome was at 5x compared to a single quantitative channel. This sample was then serially diluted in a background of carrier proteome, resulting in samples with carrier proteome levels of 20x, 50x, 100x, 250x, 500x and 1,000x. Performing the dilution in this manner ensured the relative amount of quantitative signals was constant in each experiment.

### Single cell sample preparation

HEK293 and HeLa cells were grown in Dulbecco’s modified Eagle’s medium (DMEM) supplemented with 10% fetal bovine serum, 1% MEM non-essential amino acid, 2 mM Glutamine. The incubator was operated at 37 °C and 5% CO2. Cells were harvested at 70–80% confluency.

HEK293 and HeLa cells were sorted into two types of nanoPOT chips. Single cells were sorted on nested nanoPOT N2 chips (4×4 well cluster in 3×11 array with a well diameter of 500 µm) and carrier samples were sorted on nanoPOTS chip (4×12 well array with a well diameter of 1200 µm). CellenOne system (Cellenion, France) was used for cell sorting and sample preparation. Both HEK293 and HeLa cells were stained with 10 µM Calcein AM (ThermoFisher, USA) prior to sorting.

For the single cell sample, 7 single HEK293 cells and 7 single HeLa cells were sorted in the nested nanowells per Fig. 4a. 10 nL of lysis buffer containing 0.1% DDM, 2 mM TCEP, and 5 mM CAA in 100 mM HEPES, pH 8.0 was dispensed into each nanowell. The cell sorted N2 chip was placed in a preheated humidity box and incubated at 60°C for 30 minutes, then cooled back to room temperature. Proteins were digested by adding a mixture of 0.25 ng trypsin and 0.075 ng of lys-C in 4 nL of 100 mM HEPES, pH 8.0 and incubated at 37℃ overnight in humidity condition. For TMTPro labeling, we added 10 nL (100 ng) of each of the TMTPro tags dissolved in DMSO into each of the corresponding nanowells per Fig. 4a and incubate for 1 hour at room temperature. The TMTPro labeling was quenched with 2 nL of 5% hydroxylamine to each nanowell and incubated at room temperature for 15 minutes.

For the carrier sample, we sorted an equal mixture of HEK293 and HeLa cells on the nanoPOTS chip to reach 20 (10 HeLa & 10 HEK293), 50 (25 HeLa & 25 HEK293), and 100 (50 HeLa & 50 HEK293) cells. 100 nL of lysis buffer containing 0.1% DDM, 2 mM TCEP, and 5 mM CAA in 100 mM HEPES, pH 8.0 was dispensed into each nanowell. The cell sorted nanoPOT chip was placed in a preheated humidity box and incubated at 60°C for 30 minutes, then cooled back to room temperature. Proteins were digested by adding a mixture of 0.50 ng trypsin and 0.15 ng of lys-C in 8 nL of 100 mM HEPES, pH 8.0 and incubated at 37℃ overnight in humidity condition. For TMTPro-SH labeling, we added 50 nL (500 ng) of the TMTPro-SH tag dissolved in DMSO into each of the corresponding nanowells per Fig. 4 and incubated for 1 hour at room temperature. The TMTPro labeling was quenched with 8 nL of 5% hydroxylamine to each nanowell and incubated at room temperature for 15 minutes. The single cell and its corresponding carrier samples were pooled together manually with 5 uL of 0.02 %DDM, 0.1 %FA and placed in an autosampler vial prior to LC-MS.

### Mass spectrometry acquisition

For bulk diluted samples peptides were separated using a Thermo UltiMate 3000 RSLCnano ProFlow system (Thermo Fisher Scientific) with a gradient of 4% buffer A (98% H2O, 2% ACN with 0.1% formic acid) to 30% at a flow rate of 300 nL/min over 96 min. Samples were separated over a 25-cm IonOpticks Aurora Series column (IonOpticks). Samples were analyzed by a nanoscale liquid chromatography (nLC)– MS/MS analysis of 125 min using an Orbitrap Eclipse mass spectrometer (Thermo Fisher Scientific). The native instrument method was used for MS1 and initial ion-trap MS2 analysis. MS1 survey scans were collected in the Orbitrap (200,000 resolution, AGC = 1×10^6^, max IT = 50 ms, 350-1350 m/z range) and used to select the top precursors such that an MS1 was taken every 1 sec (Top Speed selection, dynamic exclusion: select one time per 10 sec, mass tolerance = 10 ppm) using the MIPS filter, considering charge state 2-6, and using a precursor fit filter (fit error = 70% and fit window = 0.5). Precursors were isolated with a 0.5 m/z window and analyzed in the ion-trap (scan rate = Rapid, AGC = 2×10^4^, max IT = 35 ms) following fragmentation with CID at 35 NCE. Resulting ion-trap scans were analyzed further by inSeqAPI (described below).

For single cell samples peptides were separated using a gradient of 5% buffer A to 11% at 15 min (300 nL/min), 12% at 17 min (300 nL/min to 150 nL/min) to 18.5% at 40 min (150 nL/min), 32% at 114 min (150 nL/min), 45% at 126 min (150 nL/min) before column washing and equilibration. Samples were loaded onto a pre-column (Acclaim PepMap 100, 75 µm x 2 cm, C18, 3 µm and 100 Å particles, Thermo Fisher Scientific) before being separated over a 25-cm IonOpticks Aurora Series column (IonOpticks). Samples were analyzed by a nanoscale liquid chromatography (nLC)–MS/MS analysis of 140 minutes using an Orbitrap Eclipse mass spectrometer (Thermo Fisher Scientific). The native instrument method was used for MS1 and initial ion-trap MS2 analysis. MS1 survey scans were collected in the Orbitrap (200,000 resolution, AGC = 1×10^6^, max IT = 50 ms, 350-1350 m/z range) and used to select the top 15 precursors (dynamic exclusion: single charge state per precursor, select one time per 35 sec, mass tolerance = 10 ppm) using the MIPS filter, considering charge state 2-6, and using a precursor fit filter (fit error = 70% and fit window = 0.5). Precursors were isolated with a 0.5 m/z window and analyzed in the ion-trap (scan rate = Turbo, analysis mass range 200-1200 m/z, AGC = 1.5×10^4^, max IT = 254 ms) following fragmentation with CID at 35 NCE. Resulting ion-trap scans were analyzed further by inSeqAPI (described below).

### inSeqAPI acquisition methods

For SCoPE-MS FTMS2, ion-trap scans triggered by the instrument method were analyzed within inSeqAPI using a ‘Real-time Search Filter’ comprising a Comet search that was identical to the off-line Comet search, except that the database did not contain decoy entries. Following the real-time database search a ‘PPM Filter’ was utilized to adjust the systematic PPM error. Then an ‘LDA Filter’ was applied using linear-discriminant analysis (LDA) with pre-determined LDA coefficients to determine the confidence of the identification. If peptides passed this LDA filter inSeqAPI sent instructions to the mass spectrometer to perform an FTMS2 scan (isolation width - 0.5 m/z, HCD at 37.5 NCE, Orbitrap, 50,000 resolution) with variable ion sampling parameters (High - 5×10^5^ AGC and 750 ms max IT; Medium - 3.5×10^5^ AGC and 500 ms max IT; Low - 3×10^5^ AGC and 250 ms max IT), depending on the nLC-MS/MS experiment.

For SCoPE SPS-MS3, ion-trap scans triggered by the instrument were analyzed identically to SCoPE FTMS2, except that additional filters were in place following the ‘LDA Filter’. Following the ‘LDA Filter’ potential SPS ions were filtered by a ‘Mass Range Filter’ (400 - 2000 m/z), ‘Precursor Range Exclusion Filter’ (−15 m/z below, 5 m/z above precursor), ‘Tag Loss Exclusion Filter’ (tag = TMT), ‘SPS Filter’ (quant modification = ‘TMT’, tolerance = 0.35 Da), and ‘TopNFilter’ (topN = 6). If all filters were passed, an SPS-MS3 scan was sent to the instrument (CID at 35 followed by HCD at NCE 55, Orbitrap, 50,000 resolution, isolation width 1.2 m/z followed by 2 m/z, 5×10^5^ AGC, and 750 ms max IT).

For toma-scpMS FTMS2, ion-trap scans triggered by the instrument were analyzed identically to SCoPE FTMS2, except that an additional filter was in place following the ‘LDA Filter’. A ‘Modification Filter’ was added that only triggered subsequent scans when the real-time identification was of an offset carrier proteome peptide (exclude modification [−6.0138]). If all filtered were passed, inSeqAPI sent instructions to the mass spectrometer to perform an offset FTMS2 scan (isolation width - 0.5 m/z, HCD at 37.5 NCE, Orbitrap, 50,000 resolution) with variable ion sampling parameters (High - 5×10^5^ AGC and 750 ms max IT; Medium - 3.5×10^5^ AGC and 500 ms max IT; Low - 3×10^5^ AGC and 250 ms max IT), depending on the nLC-MS/MS experiment. The offset mass FTMS2 isolation m/z was determined by converting any TMT-SH modifications to TMT within the identified peptide in real-time. For single cell samples the method was identical except for the inSeqAPI triggered offset FTMS2 AGC target (3.5×10^5^), the ‘Modification Filter’ (exclude modification [−9.0239]), and that the mass offsets were determined by converting TMTpro-SH to TMTpro modifications within identified peptides in real-time.

For toma-scpMS SPS-MS3, ion-trap scans triggered by the instrument were analyzed identically to toma-scpMS FTMS2. If all filtered were passed, inSeqAPI sent instructions to the mass spectrometer to perform an offset ion-trap MS2 scan (CID at 34.5 NCE, 200-1200 m/z range, isolation width = 0.5 m/z, 1.5×10^4^ AGC, 300 ms max IT). The offset mass ion-trap isolation m/z was determined by converting any TMT-SH modifications to TMT within identified peptides in real-time. The offset ion-trap MS2 sent by inSeqAPI was then analyzed by inSeqAPI by a ‘Percent TIC Filter’ (percentTIC = 2), ‘Mass Range Filter’ (400-2000 m/z), ‘Precursor Range Exclusion Filter’ (−15 m/z below and 5 m/z above), ‘Tag Loss Exclusion Filter’ (tag = TMT), ‘SPS Filter’ (quant modification = ‘TMT’, tolerance = 0.35 Da), and ‘TopNFilter’ (topN = 6). If all filters were passed, an SPS-MS3 scan was sent to the instrument (CID at 35 followed by HCD at NCE 55, Orbitrap, 50,000 resolution, isolation width 1.2 m/z followed by 2 m/z, 5×10^5^ AGC, and 750 ms max IT). For single cell samples an identical method was used except that for select parameters in the inSeqAPI triggered ion-trap MS2 scan (max IT = 254 ms), the ‘Modification Filter’ (exclude modification [−9.0239]), the ‘Precursor Range Exclusion Filter’ (455-2000 m/z), the ‘Tag Loss Exclusion FIlter’ (tag = TMTpro), SPS-MS3 scan (CID at 34.5 NCE followed by HCD at 45 NCE, isolation width 1.2 m/z followed by 3 m/z, 7.5×10^5^ AGC), and mass offsets determined by converting TMTpro-SH to TMTpro modifications within identified peptides in real-time.

### Data analysis

Raw mass spectrometry data were converted to mzXML using msconvert within ProteoWizard version 3.0.21042 (24). The resulting mzXML files were edited prior to search to ensure that only ion-trap scans triggered by the instrument method were searched. For SCoPE-MS RTS-SPS-MS3, mzXML files were searched without modification. For SCoPE-MS RTS-FTMS2 (RETICLE) and toma-scpMS FTMS2, FTMS2 scans were edited to have a scan order of 3 within the mzXML. For toma-scpMS SPS-MS3, inSeqAPI triggered offset ion-trap MS2 scans were removed from the mzXML and the parent scan number for the offset SPS-MS3 scan was set to the instrument triggered ion-trap MS2 scan number.

Edited mzXML files were searched with Comet version 2020013 (25) against a Human database containing contaminants (UniprotKB Human database downloaded August 2017, 364,690 entries including forward and reverse proteins) (26). For the database search the precursor mass tolerance was 20 ppm around the monoisotopic mass and a fragment bin tolerance of 1.005 with a fragment bin offset of 0.4. Peptides were considered following a full in silico trypsin (with proline rule) digest allowing for one missed cleavage. For bulk diluted samples static modifications were TMT-SH on lysine and the peptide n-terminus (235.1767) as well as carbamidomethyl on cysteine (57.0215) and variable modifications were oxidized methionine (15.9949) and TMT on lysine and the peptide n-terminus (−6.0138). For single cell experiments, the static modifications were TMTpro-SH on lysine and the n-terminus (313.2310) as well as carbamidomethyl on cysteine (57.0215) and variable modifications were oxidized methionine (15.9949) and TMTpro on lysine and the peptide n-terminus (−9.0239). For all searches a maximum of 3 variable mods per peptide were considered for the database search. Resulting peptide spectral matches were filtered to 2% false discovery rate (FDR) using Percolator version 3.5 (27). A 2% threshold was chosen for offline peptide FDR because identified peptides were further filtered for cases in which an online identification occurred, the resulting FDR of quantified peptides was typically less than 1%. Results were then filtered to a protein FDR of 1% via a picked-protein approach (28). Quantitative signal-to-noise (SN) values were extracted from raw data using SCPCompanion (9) and combined with resulting identification data within R (29).

For quantitative analysis of equimolar and mixed ratio bulk diluted samples analysis was performed in R (29). For toma-scpMS equimolar ratios, interference from the carrier within the 131n channel led us to disregard this channel and continue with the analysis as a 9-plex sample. Data within quantitative channels was normalized to account for pipetting errors prior to quantitative analysis. The average signal within quantitative ‘single cell’ channels (QuantSN) was calculated by summing ‘single cell’ channels and dividing by the number of non-carrier samples within the experiment. The carrier signal was calculated by summing signals within 126 and 127c for SCoPE-MS or 131c and 132 for toma-scpMS. Carrier ratios were calculated at the PSM level by dividing the carrier signal by the average QuantSN. Measurement CVs were calculated by calculating the standard deviation of quantitative channels by the average QuantSN at the PSM. Ion injection times were also calculated at the PSM level. Quantitative cutoffs were calculated by SCPCompaion and were 126 and 142 QuantSN for SCoPE-MS and toma-scpMS equimolar mixtures, respectively as well as 63.3 QuantSN for mixed ratio samples (based on four 400 pg samples). When SCoPE-MS data was analyzed without the 128c channel the QuantSN cutoff was set to 47.25 QuantSN (based on three 400 pg samples). The number of quantified proteins reported is the number of non-decoy proteins quantified following application of protein FDR. For equimolar SCoPE-MS, the 50x carrier displayed a low number of quantified proteins that was not indicative of method performance for both FTMS2 and SPS-MS3 and these points were not plotted in Fig. 2 and Sup. Fig. 4 line plots.

For HeLa and HEK293 single cell experiments, data were subsequently analyzed in R (29). Carrier Ratio was determined by the summed signal of Carrier channels divided by the average signal within the single cell channels (average QuantSN). Peptide data were summed to the protein level before application of a summed QuantSN (total signal within single cell channels) cutoff of 220. Reported quantified proteins represent non-decoy proteins following the application of protein FDR. Data from multiple plexes were normalized per protein across all channels before being combined. PCA was performed within R on normalized and combined data. Log2 ratios were calculated using all HeLa and HEK293 samples across all experiments. Subsequently, p-values were calculated using a Student’s t-test (two tails, unequal variance) and adjusted by Benjamini-Hochberg. Protein changes were called significant if the adjusted p-value was ≤ 0.05.

## Results

### Offset mass carrier proteome for single cell proteomics

Multiplexed scpMS approaches (e.g., SCoPE-MS or iBASIL) utilize a 20-500x carrier proteome to enable the detection of precursor peptides within MS1 survey scans and improve the identification of peptides by contributing abundant fragment ions within identification scans (**Fig. 1a**). While a carrier proteome improves the number of peptides identified within a scpMS experiment, the presence of highly-abundant carrier proteome reporter ions may reduce the quantitative accuracy of single cell proteome measurements (9). This effect can be somewhat overcome by increasing ion sampling, however both the carrier and single cell precursor ions are isolated together such that sampling more single cell proteome ions requires additional sampling of carrier proteome ions (**Fig. 1a**).

**Figure 1.**
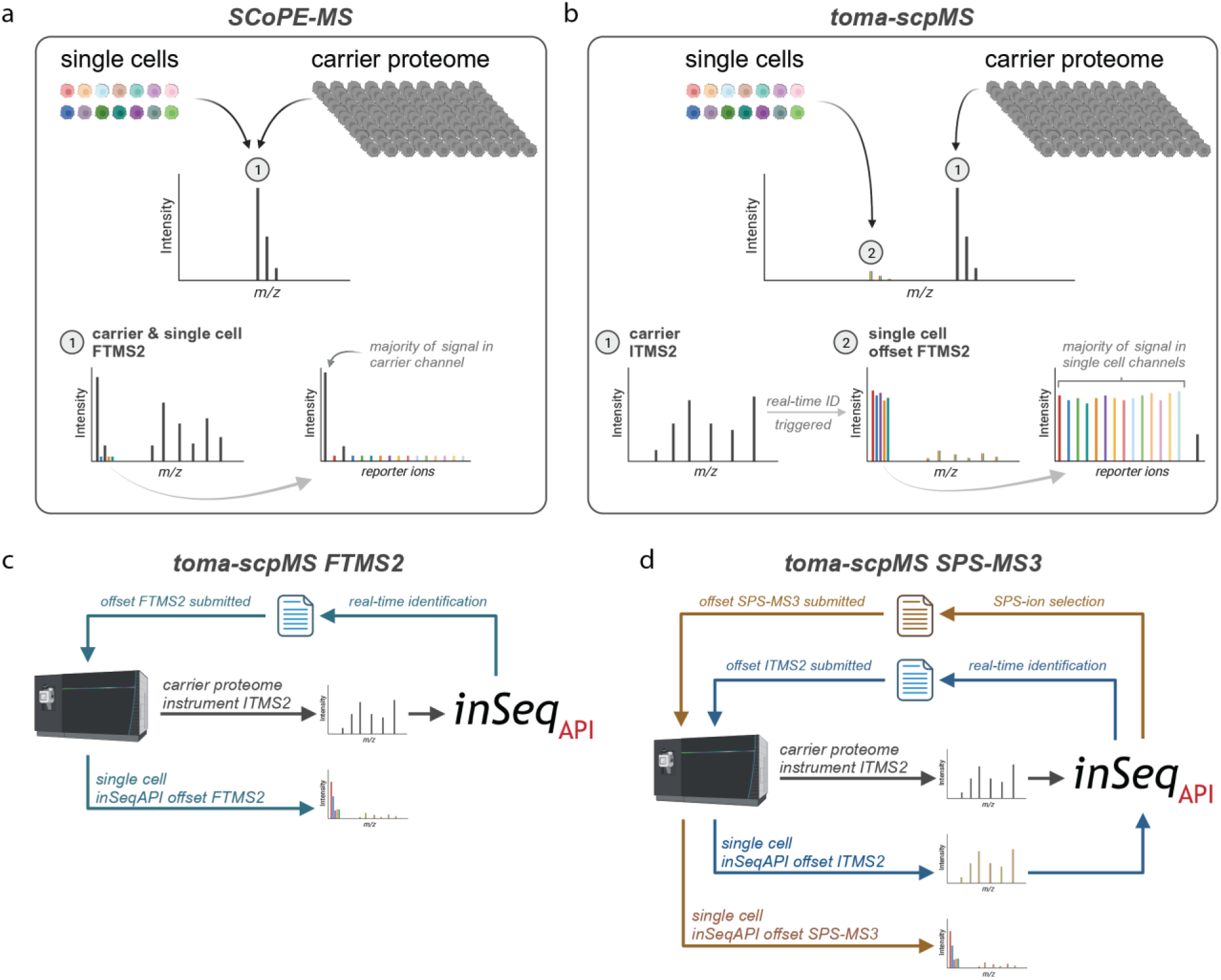
SCoPE-MS and toma-scpMS schemes and toma-scpMS inSeqAPI workflow. **a**, Diagram of single cell proteomics by mass spectrometry (SCoPE-MS) workflow. **b**, Diagram of the targeted by offset mass acquisition single cell proteomics (toma-scpMS) workflow. **c**, Workflow for toma-scpMS FTMS2 inSeqAPI method. **d**, Workflow for toma-scpMS SPS-MS3 inSeqAPI method.

We surmised that separating the carrier and single cell proteomes would maintain the benefits of the carrier proteome for peptide identification while also enabling increased sampling of single cell peptides to improve quantification (**Fig. 1b**). In principle, this approach is similar to the targeted peptide and sample multiplexing approach TOMAHAQ (30, 31), however we sought to employ this method in an untargeted, discovery workflow. To accomplish this we utilized the instrument application programming interface (iAPI) to create an intelligent data acquisition (IDA) program called inSeqAPI that enables a real-time Comet (18, 25) database search and subsequent instrument control. Our program, inSeqAPI, is similar to other iAPI based IDA implementations (17–19), but enables creation of instrument methods similar to the native instrument method editor with methods consisting of one or more ‘Nodes’ which comprise ‘Filter Groups’, ‘Filters’, and ‘Scans’ (**Sup. Fig. 1a-b**). This structure enables the recapitulation of methods that are built within the native method editor as well as the construction of complex acquisition methods that can not be typically performed.

Here, we constructed a triggered by <u>o</u>ffset <u>m</u>ass <u>a</u>cquisition method for scpMS (toma-scpMS) that utilized the native instrument method to perform ion trap MS2 (ITMS2) scans and subsequently captured the resulting data for analysis within inSeqAPI (**Fig. 1c-d**). MS2 spectra were searched in real-time by Comet against a database comprised of both carrier and single cell proteome peptides and identification of a carrier proteome peptide prompted inSeqAPI to calculate the m/z of the partner single cell peptide and instruct the mass spectrometer to perform a quantitative scan of the corresponding single cell peptide. Quantitative scans followed either an FTMS2 (**Fig. 1c**) or SPS-MS3 workflow (**Fig. 1d**). For FTMS2 (toma-scpMS FTMS2), inSeqAPI instructed the instrument to collect an offset FTMS2 to quantify the offset single cell peptide (**Fig. 1c, Sup. Fig 1a**). For SPS-MS3 (toma-scpMS SPS-MS3), inSeqAPI prompted the collection of an offset ITMS2 scan on the single cell peptide, this scan was then analyzed by inSeqAPI to select SPS ions for a subsequent single cell SPS-MS3 scan sent by inSeqAPI (**Fig. 1d, Sup. Fig 1b**).

### Evaluation of an offset mass carrier proteome for equimolar mixtures

Due to the separate isolation of the carrier and single cell peptides an offset mass carrier proteome should enable the utilization of higher carrier proteome levels. Due to this, we sought to compare FTMS2 and SPS-MS quantification for toma-scpMS and SCoPE-MS at carrier proteome levels up to 1000x. To analyze these samples four instrument methods were implemented within inSeqAPI. Two of these methods were used to analyze SCoPE-MS samples: 1) ‘RTS-FTMS2’ (RETICLE) (22), an ion trap scan followed by a real-time peptide identification triggered FTMS2 (**Sup. Fig. 2a**); and 2) ‘RTS-SPS-MS3’ an ion trap scan followed by a real-time identification triggered SPS-MS3 (**Sup. Fig. 2b**). The two methods described in Fig. 1 were used to analyze samples containing an offset mass carrier proteome: 1) ‘toma-scpMS FTMS2’, an ion trap scan of a carrier peptide followed by a real-time identification triggered FTMS2 on the single cell peptide (**Sup. Fig. 2c**); and 2) ‘toma-scpMS SPS-MS3’, an ion trap scan of a carrier peptide followed by a real-time identification triggered ion trap scan of a single cell peptide which was used to select SPS ions for a subsequent SPS-MS3 scan of the single cell peptide (**Sup. Fig. 2d**)

We utilized these methods to analyze separate mixtures of bulk-diluted peptides simulating ‘single cell’ SCoPE-MS and toma-scpMS samples with carrier levels of 5x, 20x, 50x, 100x, 250x, 500x, and 1000x (**Fig. 2a**). For SCoPE-MS, we utilized TMT to create an 8-plex sample with the carrier in channel 126, an empty 127c channel, and the remaining eight channels filled with 400 pg of a tryptic K562 digest (**Fig. 2a**). For toma-scpMS we utilized TMT and TMT-SH to create a 10-plex sample with 400 pg of tryptic K562 digest in channels 126-131n and the carrier labeled with TMT-SH (**Fig. 2a**). Note, the TMT-SH label produces a 132 reporter ion, theoretically enabling more channels to be utilized in the toma-scpMS approach. However, subsequent analysis of the reporter ion intensities demonstrated that within toma-scpMS FTMS2 data the 131n channel experienced noticeable interference from the −1 Da impurity of the TMT-SH reporter ion beginning at 100x carrier proteome (**Sup. Fig. 3a-d**). This was due to the 132 reporter ion containing both the ‘n’ and ‘c’ heavy atoms and producing interference into both the 131n and 131c channels. As a result, the 131n channel was disregarded and toma-scpMS data were analyzed further as a 9-plex experiment.

**Figure 2.**
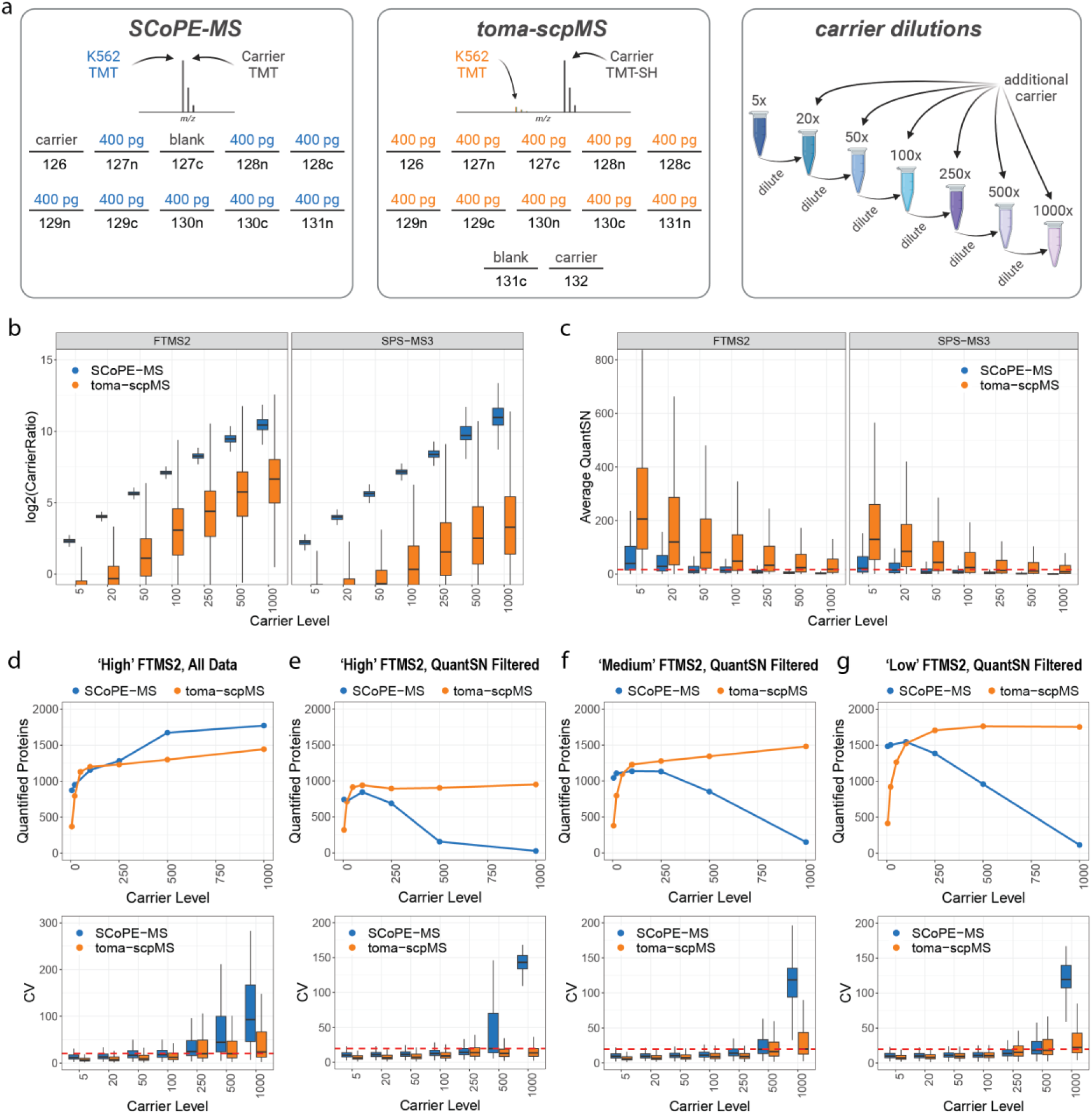
Comparison of SCoPE-MS and toma-scpMS for the analysis of equimolar mixtures. **a**, Schematics describing the sample composition and dilution scheme for the creation of equimolar bulk-diluted ‘single cell’ mixtures. **b**, Log2 transformation of the measured carrier ratio for SCoPE-MS and toma-scpMS as calculated by the sum of carrier reporter ion channels divided by the average of ‘single cell’ channels. **c**, Average QuantSN (summed signal-to-noise ratio across ‘single cell’ channels divided by the number of ‘single cell’ channels) for SCoPE-MS and toma-scpMS. The dotted line represents a QuantSN threshold of 16, the approximate threshold to be considered quantified. **d**, The number of quantified proteins and Coefficient of variation (CV) when all data is considered for toma-scpMS FTMS2 using ‘High’ ion sampling parameters. **e-g**, CV and quantified proteins for QuantSN filtered data collected using ‘High’, ‘Medium’, or ‘Low’ ion sampling parameters, respectively. For all box plots, the line denotes the median while the box denotes the interquartile range and the whiskers the highest and lowest value, excluding outliers (not shown).

As described above, the offset mass carrier proteome enables isolation of single cell peptides separate from the matching peptide within the carrier proteome. However, interfering peptides, with different sequences than the single cell peptide, may be co-isolated during quantitative analysis and result in carrier proteome reporter ion within the quantitative scan of toma-scpMS methods. Previously, we have described how SCPCompanion measures the observed carrier proteome ratio by dividing the sum of the signals for the carrier reporter ions (including impurities) by the average signal within single cell reporter channels (9). To characterize the level of carrier proteome reporter ions measured from co-isolated carrier proteome (SCoPE-MS) or interfering peptides (toma-scpMS) interference, we calculated the carrier proteome ratio within each quantitative scan for all analyses (**Fig 2b**). We found the measured carrier proteome ratio was dramatically lower for toma-scpMS as compared to SCoPE-MS. For example, within the 1000x carrier sample SCoPE-MS exhibited carrier ratios of ∼1400x and ∼1900x for FTMS2 and SPS-MS3, while toma-scpMS data exhibited carrier ratios of ∼100x and ∼10x for FTMS2 and SPS-MS3, respectively (**Fig 2b**). These data demonstrated that the offset carrier proteome approach was able to decrease the amount of carrier proteome ion by 14x for FTMS2 and 190x for SPS-MS3 analysis.

Based on the reduction of the co-isolated carrier proteome we concluded that the majority of the peptide ions isolated within toma-scpMS were ‘single cell’ ions. This was demonstrated by calculating the average signal-to-noise per ‘single cell’ channel (QuantSN) by summing the ‘single cell’ reporter ion signals and dividing the result by the number of multiplexed samples (**Fig. 2c**). We found that average QuantSN generally decreased as the carrier level increased, but toma-scpMS methods generally produced ∼5x more QuantSN at a given carrier level (**Fig. 2c**). Furthermore, for toma-scMS FTMS2 the lower bound of the interquartile range remained above the QuantSN cutoff of 16 SN/channel (red line, **Fig. 2c**). For both SPS-MS3 methods the average QuantSN was consistently lower than the matching FTMS2 method and for SCoPE-MS only the 5x and 10x carrier samples exhibited a median QuantSN value at or above the quantification threshold (**Fig. 2c**). Conversely, toma-scpMS methods maintained a median QuantSN at or above the threshold until the carrier level reached 250x (**Fig. 2c**).

When all quantitative data were considered, increased carrier proteome levels led to a larger number of protein identifications (**Fig. 2d, Sup. Fig. 4a -‘All Data’**), particularly for SCoPE-MS. This was due to a dramatic reduction in SCoPE-MS ion injection times for quantification scans within 100-100x carrier proteome samples (**Sup. Fig 4b**). The decreased injection time was due to the ion injection time calculation being dependent on the precursor peptide peak which is a combination of both the carrier and single cell proteomes within SCoPE-MS. For toma-scpMS, injection times are dependent on the level of the multiplexed single cell proteomes and appeared to be stable until 500x carrier, at which point they declined (**Sup. Fig 4b**). This decline was attributed to precursor interference within the offset single cell proteome isolation window causing a decrease in the predicted injection time required to reach an ion sampling target.

Previously, we have demonstrated that post-analysis filtering based on signal-to-noise was a key component of controlling measurement variability (9). Using SCPCompanion we determined a QuantSN cutoff of 126 for the 8-plex SCoPE-MS samples and 142 for the 9-plex toma-scpMS samples (**Fig. 2e, Sup. Fig. 4a -‘QuantSN Filtered’**) (9). After applying this cutoff we found that SCoPE-MS produced more quantified proteins at carrier levels below 100x (**Fig. 2e, Sup. Fig. 4a**). This was attributed to increased spectral complexity of toma-scpMS where each peptide was represented by two precursor clusters (one each from the carrier and single cell proteomes) within MS1 survey scans. However, at carrier levels 100x and above the offset carrier proteome became the dominant species within the survey scans and the number of proteins quantified by toma-scpMS matched and then surpassed SCoPE-MS, reaching a plateau ∼900 proteins for FTMS2 (**Fig. 2e**) and ∼620 proteins for SPS-MS3 (**Sup. Fig. 4a**). Conversely, SCoPE-MS experienced a dramatic decrease in quantified proteins above 100x carrier proteome (**Fig. 2e, Sup. Fig. 4a**). This was a result of the previously described challenge of sampling sufficient ‘single cell’ channels in the presence of high levels of carrier proteome (9). The measurement variability was then assessed by plotting the coefficient of variation (CV) for the quantified proteins in each analysis method (**Fig. 2e, Sup. Fig. 4c**). Here we found that toma-scpMS consistently produced measurements with lower CVs as compared to SCoPE-MS (**Fig. 2e, Sup. Fig. 4c**). This result is a direct result of the higher QuantSN demonstrated by toma-scpMS methods because higher intensity measurements lead to lower variability of quantification (9).

Previous efforts to optimize single cell proteomics data depth and quantitation quality have focused on increased ion sampling through adjustment of the automated gain control (AGC) and maximum injection time (max IT) (4, 9, 12). The data above relates to ‘High’ ion sampling conditions (5×10^5^ AGC, 750 ms max IT), but we also tested ‘Medium’ (3.5×10^5^ AGC, 500 ms max IT) and ‘Low’ (3×10^5^ AGC, 250 ms max IT) ion sampling conditions (**Fig. 2f-g**). Decreasing ion sampling conditions led to an overall increase in quantified proteins, however the measurement variability of toma-scpMS and SCoPE-MS remained similar to the ‘High’ sampling conditions. At low carrier levels SCoPE-MS quantifies more proteins and at 100x carrier proteome the methods are equal. However, when the carrier proteome is >100x toma-scpMS quantifies more proteins with ∼1,500 and ∼1,700 proteins for ‘Medium’ and ‘Low’ ion sampling, respectively (**Fig. 2f-g**). Analysis of measurement variability demonstrated that toma-scpMS utilizing ‘Medium’ ion sampling consistently produced measurements with lower CVs as compared to SCoPE-MS (**Fig. 2f**), but the differences in CV with ‘Low’ ion sampling parameters were negligible (**Fig. 2g**). Furthermore, for carrier levels above 500x for Medium and 250x for Low ion sampling toma-scpMS quantification became more variable such that the upper bound the interquartile range is above 20% CV (**Fig. 2f-g**). This demonstrates that ion sampling parameters must still be carefully considered, even when employing an offset mass carrier proteome.

### Evaluation of an offset mass carrier proteome for mixed ratios

Equimolar mixtures enable characterization of measurement variability, but accuracy is best measured with a mixed ratio sample. To understand the utilization of an offset mass carrier proteome for the quantification of mixed ratios we created 8-plex 1:2 mixtures for both SCoPE-MS and toma-scpMS where 1x and 2x channels contained 400 pg and 800 pg on column, respectively (**Fig. 3a**). We combined this mixture with carrier proteome to produce samples with 5x, 20x, 50x, 100x, 250x, 500x, and 1000x carrier proteome (**Fig. 3a**). These samples were analyzed with the previously described methods (**Sup. Fig. 2**) and the resulting data were filtered with a QuantSN ≥126 SN. Similar to 1:1 mixtures, SCoPE-MS outperforms toma-scpMS for low carrier levels (5x-50x), however at high carrier proteome levels (250x-1000x) toma-scpMS quantifies more proteins as compared to SCoPE-MS (**Fig. 3b**).

**Figure 3.**
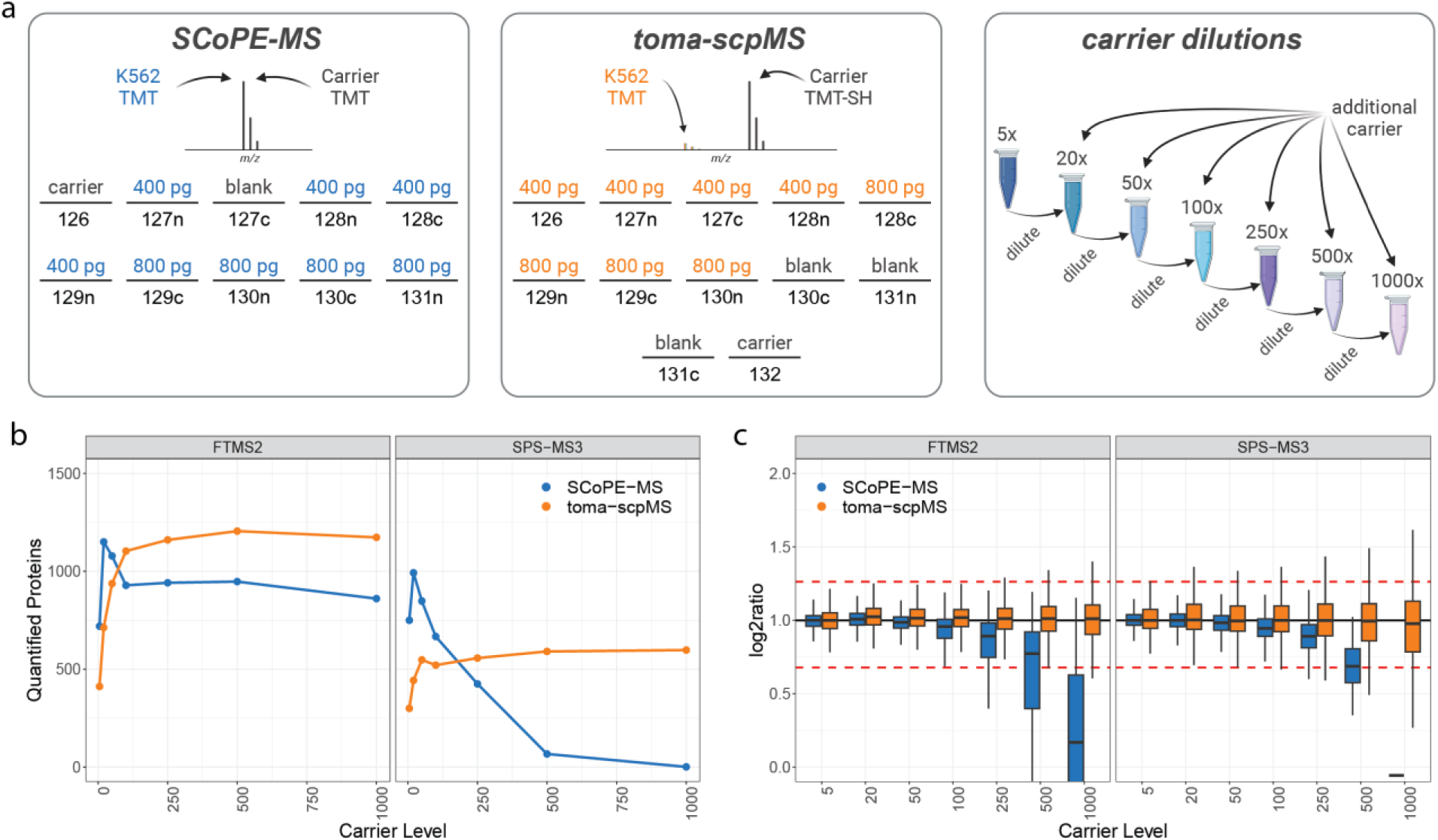
Comparison of SCoPE-MS and toma-scpMS for analysis of mixed ratio samples. **a**, Schematics describing the sample composition and dilution scheme for the creation of mixed ratio bulk-diluted ‘single cell’ mixtures. **b**, The number of quantified proteins after applying a QuantSN cutoff at the peptide spectral match (PSM) level. **c**, Distribution of measured protein ratios for the mixed ratio. The target ratio was 2 (log2ratio = 1, black horizontal line) and the red dotted lines represent a 20% measurement error. For box plots, the line denotes the median while the box denotes the interquartile range and the whiskers the highest and lowest value, excluding outliers (not shown).

We then calculated the fold change for each quantified protein to determine the quantitative accuracy of SCoPE-MS and toma-scpMS (**Fig. 3c**). Curiously, we found that SCoPE-MS demonstrated noticeable ratio compression with as little as 100x carrier proteome (**Fig. 3c**). This was particularly interesting because these samples did not contain a background interference proteome, demonstrating that ratio compression will occur at high carrier proteome levels independent of background interference. Ratio compression was not detected for toma-scpMS FTMS at any carrier proteome level, but a slight compression was present for the 1000x carrier for toma-scpMS SPS-MS3 (**Fig. 3c**). There was a slight increase in the median ratio for toma-scpMS FTMS2 that may have been due to interference from the −2 Da impurity of the carrier proteome label within the 130n and 130c reporter ion channels (**Fig. 3c**). This is supported by the elimination of this ratio increase for toma-scpMS SPS-MS3, where the amount of carrier proteome measured in the quantification scan is lower than toma-scpMS FTMS2 (**Sup. Fig. 5**). Together these data demonstrate that toma-scpMS provides deeper proteome coverage and more accurate quantification data at higher carrier levels as compared to SCoPE-MS.

### Single cell analysis utilizing an offset mass carrier proteome

Having demonstrated the advantages of the toma-scpMS approach on diluted bulk samples, we next employed toma-scpMS FTMS2 and SPS-MS3 for the analysis of single cells (**Fig. 4**). HeLa and HEK293 cells were analyzed by multiplexed experiments containing 14 single cells (7 HeLa and 7 HEK293) and either 20x, 50x, or 100x offset carrier proteome (**Fig. 4a**). Sample preparation was performed using a CellenOne system to sort single cells into either an N2 (32) or nanoPOTS (33) chip for the multiplexed single cell or the carrier proteome preparation, respectively (**Fig. 4a**). For the carrier proteome 20, 50, or 100 cells (equal mixture of HeLa and HEK293) were prepared alongside single cells to better match the single cell and carrier proteomes. Cells were reduced, alkylated, digested with trypsin/lysC, and labeled with either the appropriate TMTpro channel for single cells or TMTpro-SH for carrier proteomes (**Fig. 4a**). Following sample preparation, multiplexed single cells were combined with carrier proteome and analyzed with either a toma-scpMS FTMS2 (three plexes totaling 42 single cells per carrier level) or SPS-MS3 method (one plex totaling 14 single cells per carrier level) (**Fig. 4a**). Data were collected with 3.0×10^5^ AGC, 750 ms max IT ion sampling parameters.

**Figure 4.**
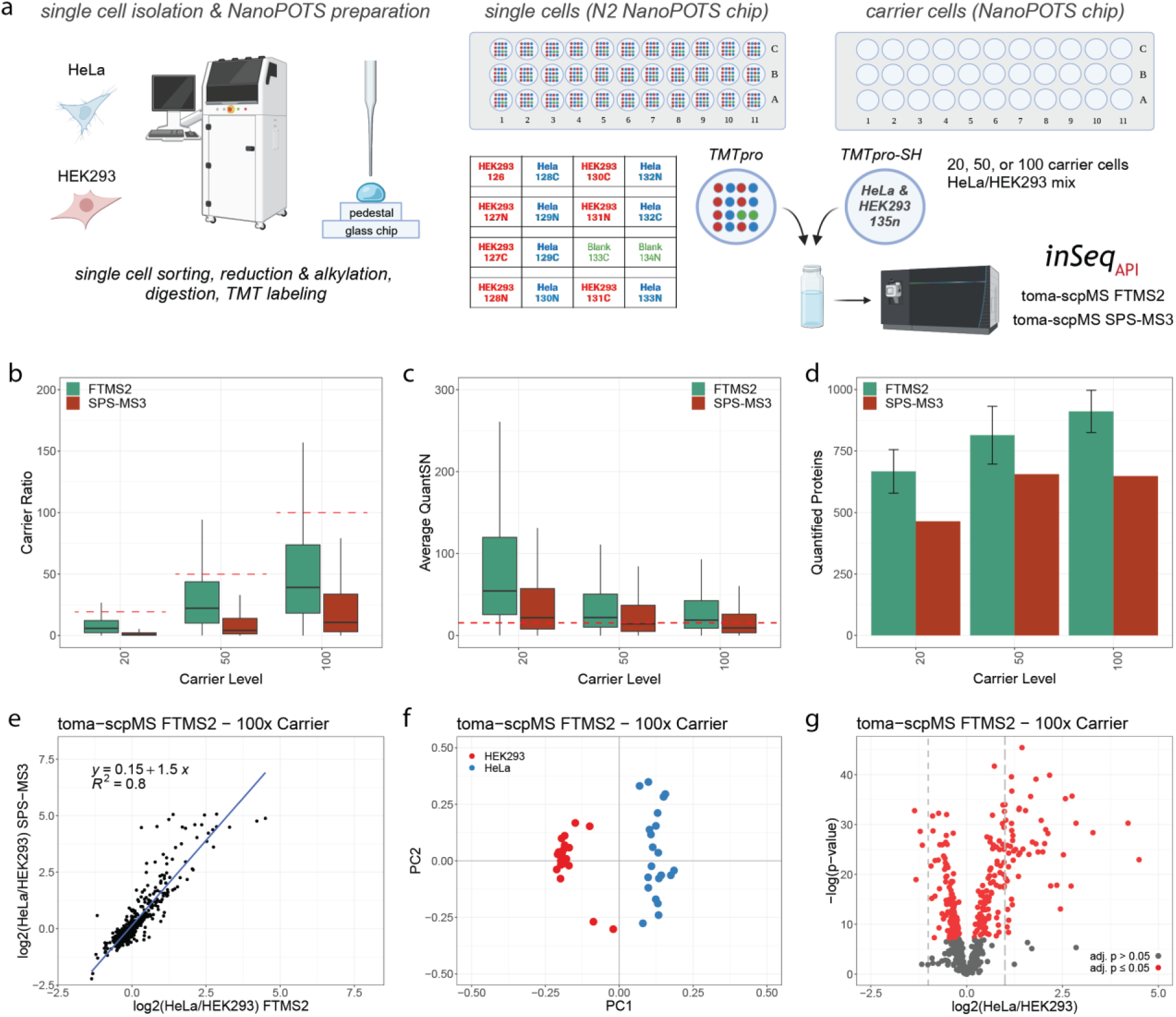
Analysis of single cells from two cell types with toma-scpMS. **A**, Schematic describing the preparation of single cells for analysis by toma-scpMS. Cells were isolated and prepared by the CellenOne instrument using either an N2 NanoPOTS chip (single cells) or a NanoPOTS chip (carrier proteome). The single cells and carrier cells were labeled as described for all multiplexed experiments. Three plexes were analyzed with toma-scpMS FTMS2 for the 20x, 50x, and 100x carrier proteome levels, while one plex was analyzed for each carrier level for toma-scpMS SPS-MS3. **b**, The measured carrier ratio as calculated by the sum of carrier reporter ion channels divided by the average of single cell channels. **c**, Average QuantSN (summed signal-to-noise ratio across single cell channels divided by the number of ‘single cell’ channels). The dotted line represents a QuantSN threshold of 16, the approximate threshold to be considered quantified. For all box plots in c-d, the black line denotes the median while the box denotes the interquartile range and the whiskers the highest and lowest value, excluding outliers (not shown). **d**, The number of quantified proteins for each carrier level analyzed by toma-scpMS FTMS2 or SPS-MS3. For FTMS2 bars represent the average number of proteins quantified across three plexes and the bars represent +/− SD. **e**, Correlation of toma-scpMS FTMS2 and SPS-MS3 measurements for proteins quantified in both methods. **f**, Principal component analysis based on proteins quantified across all three plexes differentiates cell types. **g**, Volcano plot displaying the differential analysis of protein abundance across the two cell types.

As described above, an offset carrier proteome will lead to the co-isolation of single cell and carrier proteome peptides of different sequences. To determine the level of co-isolated interference we calculated the carrier proteome ratio for each sample and toma-scpMS method (**Fig. 4b**). For toma-scpMS FTMS2 we found that the measured carrier proteome level was generally ≤50% of the target carrier proteome level, measuring 6x (20x), 22x (50x), and 39x (100x) (**Fig. 4b**). For toma-scpMS SPS-MS3 the estimated carrier proteome ratio was decreased as much as 10-fold, measuring 1x (20x), 4x (50x), and 10x (100x) (**Fig. 4b**). By decreasing the amount of carrier proteome reporter ions, we increase the dynamic range of single cell ion measurements and enable more accurate quantification, as supported by the diluted bulk mixed ratio samples above.

While the dramatic reduction of the carrier proteome reporter ions appears to favor quantitation by toma-scpMS SPS-MS3, the gas phase purification leads to a reduction of QuantSN. To characterize the level of QuantSN for both methods the summed QuantSN was extracted for all PSMs within each sample and data collection method. We found a general decrease in QuantSN as the level of carrier proteome increases, matching the results of our diluted bulk samples above (**Fig. 4c**). For toma-scpMS FTMS2, a large proportion of PSM measurements were above the average QuantSN cutoff of ∼16 SN calculated by SCPCompainon for all carrier levels (**red line, Fig. 4c**) This was only the case for the 20x carrier sample analyzed by toma-scpMS SPS-MS3, as ∼50% of PSMs failed to acquire more than 220 SN for the 50x and 100x carrier samples (**Fig. 4c**). After applying the QuantSN filter, we found that toma-scpMS FTMS on average quantified 667, 814, and 911 proteins for the 20x, 50x, and 100x carrier proteome samples, respectively (**Fig. 4d**). The number of quantified proteins per plex was lower with toma-scpMS SPS-MS3 as 464, 656, and 648 proteins were quantified in the 20x, 50x, and 100x carrier proteome samples, respectively (**Fig. 4d**). Interestingly, it appears that the number of proteins quantified by toma-scpMS FTMS2 increased from 50x to 100x carrier proteome, while the number of quantified proteins remained roughly the same for toma-scpMS SPS-MS3 (**Fig. 4d**). This was likely due to the lower QuantSN observed for toma-scpMS SPS-MS3 scans leading to more data being filtered out due to falling below the quantitative threshold.

While toma-scpMS SPS-MS3 quantified fewer proteins, it was possible that the gas phase purification improved quantification accuracy such that the dynamic range of measurements was larger. To understand the potential impact of ratio compression within toma-scpMS we compared the log2(HeLa/HEK293) ratio for proteins in toma-scpMS FTMS2 as compared to SPS-MS3 for each carrier proteome level (**Fig. 4e, Sup. Fig. 6a-b**). The slope of a calculated regression line can be used to characterize the level of ratio compression as values larger than one relate to higher ratios measured with the SPS-MS3 approach. Surprisingly, we found a moderate impact of ratio compression as the slope of the regression lines were 1.3, 1, and 1.5 for the 20x, 50x, and 100x carrier levels, respectively (**Fig. 4e, Sup. Fig. 6a-b**). Note, the low level of ratio compression may have been due to these data representing high abundance proteins that are less impacted by precursor interference. Nonetheless, these results are encouraging because SPS-MS3 analysis dramatically decreases QuantSN for single cell measurements, making toma-scpMS FTMS2 the preferred method.

Lastly, we sought to demonstrate that toma-scpMS FTMS2 data is sufficient to separate cell types and determine cell-type specific proteins. To demonstrate a separation of cell types we performed PCA and found that each toma-scpMS approach easily separated cell types (**Fig. 4f, Sup. Fig. 6c-g**). Interestingly, HeLa cells demonstrated higher variability as compared to HEK293 cells across all experiments, we attribute this to variations in HeLa cell sizes as there were no clear associations with TMT-plex or channel (**Fig. 4f, Sup. Fig. 6c-g**). We calculated the log2(HeLa/HEK293) ratios and determined significantly changing proteins (P<0.05, student’s t-test, two tails, unequal variance, Benjimini-Hochberg correction) (**Fig. 4g, Sup. Fig. 6h-i**). For FTMS2 we found a similar number of significantly changing proteins (20x: 212, 50x: 290, 100x: 260) as well as significantly changing proteins with levels altered more than 2-fold (20x 68, 50x: 68, and 100x: 64) (**Fig. 4g, Sup. Fig. 6h-i**). These results demonstrate that a similar number of significantly changing proteins can be detected irrespective of carrier proteome level when employing toma-scpMS FTMS2.

## Discussion

Single cell proteomics by mass spectrometry (scpMS) was greatly accelerated by SCoPE-MS, the first single cell analysis of cancer cell lines (10). An enabling feature of the initial SCoPE-MS approach was the carrier proteome - a sample containing 200 cells that was added to improve peptide selection and identification (10). Since this description, the carrier proteome has been extensively studied and the field has reached a consensus on the importance of optimizing ion sampling (AGC and max IT) for producing low-variability, high accuracy data. However, the level of carrier proteome used within scpMS remains variable with as little as 20x or as much as 500x carrier proteome being employed. The continued reliance on higher carrier proteome levels, suggests that an approach that is more robust to high carrier proteome levels would be important for scpMS analyses.

Here we introduce a triggered by offset mass acquisition method for scpMS (toma-scpMS) that utilizes an offset carrier proteome that is more robust to the utilization of higher carrier proteome levels as compared to SCoPE-MS. To enable data acquisition of samples utilizing an offset mass carrier proteome we constructed an intelligent data acquisition (IDA) method within inSeqAPI, an instrument application interface (iAPI) program that enables real-time analysis of data and instrument control. Four inSeqAPI methods were constructed to analyze toma-scpMS or SCoPE-MS samples by FTMS2 or SPS-MS3. While the SCoPE-MS methods implemented within inSeqAPI can be reproduced through the native method editor, the toma-scpMS methods are only possible when utilizing the iAPI. This is because implementing toma-scpMS required the conversion of an identified carrier proteome peptide to the equivalent single cell proteome peptide. To accomplish this, identified peptides were reduced to their unmodified form and modifications corresponding to the single cell proteome were added to the base peptide sequence. This ensured the appropriate precursor, fragment, and potential SPS ions were calculated for toma-scpMS FTMS or SPS-MS3 analysis. Currently this logic is not available within the native method editor.

Through the analysis of equimolar diluted bulk samples, we compared measurement variability for SCoPE-MS and toma-scpMS in the presence of carrier levels ranging from 5x to 1000x. These data demonstrated that toma-scpMS produced higher levels of reporter ion signal within ‘single cell’ channels (QuantSN), leading to lower variability measurements across all carrier proteome levels. Interestingly, we noted that the number of quantified proteins was higher for SCoPE-MS at the 5x and 20x carrier proteome levels. This was likely due to a greater spectral complexity for the toma-scpMS samples as both the single cell and carrier proteome peptides are readily detectable within an MS1 survey scan at low carrier levels. The two methods appeared to quantify an equal number of proteins at 100x carrier proteome level, before the number of quantified proteins dropped precipitously for SCoPE-MS. Conversely, toma-scpMS maintained the number of quantified proteins as the carrier level increased, demonstrating a greater robustness to the level of carrier proteome employed.

By analyzing a diluted bulk 1:2 mixed ratio sample we were able to compare the accuracy of toma-scpMS and SCoPE-MS. While the trend in quantified proteins was similar to that detected in the equimolar mixtures, the analysis of the ratios revealed ratio compression within SCoPE-MS beginning at 50x carrier proteome. Interestingly, these samples did not contain an equimolar interference proteome, so the ratio compression could not be due to precursor co-isolation. We attribute the ratio compression within SCoPE-ms to a truncation of the ‘single cell’ reporter ion channels due to a highly abundant carrier proteome reporter ion channel. We previously reported this effect and demonstrated how the removal of the carrier proteome reporter ion can increase the dynamic range of measured ‘single cell’ signals (9). This appears to be a fundamental limitation of utilizing high levels of carrier proteomes within SCoPE-MS. Unlike SCoPE-MS, toma-scpMS did not experience ratio compression within the mixed ratio sample, rather we observed ratio expansion with high levels of carrier proteome and toma-scpMS FTMS analysis. We surmised this was due to −2 Da interference from co-isolated carrier proteome reporter ion interfering with the ‘single cell’ signals in the 130n and 130c channels. This was supported by the elimination of this effect for toma-scpMS SPS-MS3 where interfering co-isolated peptide signal was drastically reduced. Together these data strongly support the utilization of toma-scpMS when utilizing carrier levels ≥100x.

Through the analysis of single cells we demonstrated that toma-scpMS can produce high-quality data using either an FTMS2 or SPS-MS3 quantification strategy. Within these single cell samples we demonstrate that the measured carrier proteome ratio is less than 50% the amount within the sample for toma-scpMS FTMS and less than 10% for toma-scpMS SPS-MS3, decreasing the negative impact of highly abundant carrier reporter ions. These data also demonstrated that increased carrier proteome levels reduced QuantSN within single cell channels such that the number of proteins plateaued for SPS-MS3 quantification. The number of quantified proteins with FTMS2 increased from 50x to 100x carrier proteome, however the percentage of measurements above the quantitative threshold fell. This suggests that increasing the carrier proteome further would require a concomitant increase in ion sampling which would likely reduce quantified proteins due to longer duty cycle. Encouragingly, when comparing log2(HeLa/HEK293) ratios for toma-scpMS FTMS2 and SPS-MS3 analyses we found only minor ratio compression. From these data we conclude that FTMS2 data acquisition remains superior to SPS-MS3 due to higher reporter ion signal and that 100x carrier proteome would be the recommended limit for offset carrier proteome within toma-scpMS. Lastly, we demonstrate that toma-scpMS FTMS2 acquisition produces data that enables separation of cell types and the identification of cell-type specific proteins. These results demonstrated larger variability in HeLa cells as compared to HEK293 cells, however this variation could not be associated with TMT-plex or label and was likely due to differences in HeLa cell size. This was supported by observations of cell size variance during cell sorting.

Previous exploration of using an offset carrier proteome utilized the offset scan for both identification and quantification, but found that the number of identified peptides and proteins decreased substantially as carrier levels increased (14). An important consideration for toma-scpMS is that identifications are determined by carrier proteome scans and single cell reporter ion signals are extracted from offset quantitative scans. One concern with this approach is that single cell reporter ions measured in an offset scan are not derived from the identified carrier proteome peptide. We note here that this is also a concern for SCoPE-MS, as reporter ion signals within single cell channels may be derived from co-isolated interference. For SCoPE-MS the carrier proteome fragment ions mask detection of single cell peptide fragment ions that would demonstrate detectable presence of single cell ions. From our data, the correlation of toma-scpMS FTMS2 and SPS-MS3 ratios demonstrates that quantification within toma-scpMS is faithful to the identified peptides. This is due to SPS-MS3 data being highly specific as a result of isolating the single cell precursor and then subsequently single cell fragment ions before quantification.

Overall, the data presented support the implementation of a toma-scpMS experiment utilizing an offset mass carrier proteome and FTMS2 quantification. This approach will quantify a similar number of proteins as SCoPE-MS, but produce higher quality measurements that are less impacted by ratio compression caused by the presence of high amounts of carrier reporter ions. Furthermore, toma-scpMS is more robust to high levels of carrier proteome and could be valuable in cases where the level of carrier proteome added to a scpMS experiment can not be carefully controlled.

## Abbreviations

MS: mass spectrometry
scpMS: single cell proteomics by mass spectrometry
toma-scpMS: triggered by offset mass acquisition scpMS
FTMS2: Orbitrap MS/MS scan
SPS: synchronous precursor selection
SPS-MS3: synchronous precursor selection with Orbitrap MS3 scan
SCoPE-MS: single cell proteomics by mass spectrometry
DDA: data dependent acquisition
DIA: data independent acquisition
mDIA or plexDIA: multiplexed DIA
iAPI: instrument application programming interface
CV: coefficient of variation
PSM: peptide spectral match
ms: millisecond
iBASIL: improved boosting to amplify signal with isobaric labeling

## Data Availability

Raw data have been deposited in MASSIVE (34) with the accession MSV000095629. The inSeqAPI program is available upon request and requires an instrument API agreement with ThermoFisher.

## Author Contribution Statement

T.K.C, Y.Z., and C.M.R. performed experiments. C.M.R. conceived of project, led experiments, and wrote manuscript.

## Conflicts of interest

T.K.C., Y.Z., and C.M.R. are employees of Genentech, Inc. and shareholders of Roche.

## Acknowledgements

We thank Sarah Williams for help with the nanoPOTS chip as well as members of the Proteomic and Genomic Technologies department at Genentech for helpful discussions during the writing of the manuscript.

---

**Supplementary Figure 1.**
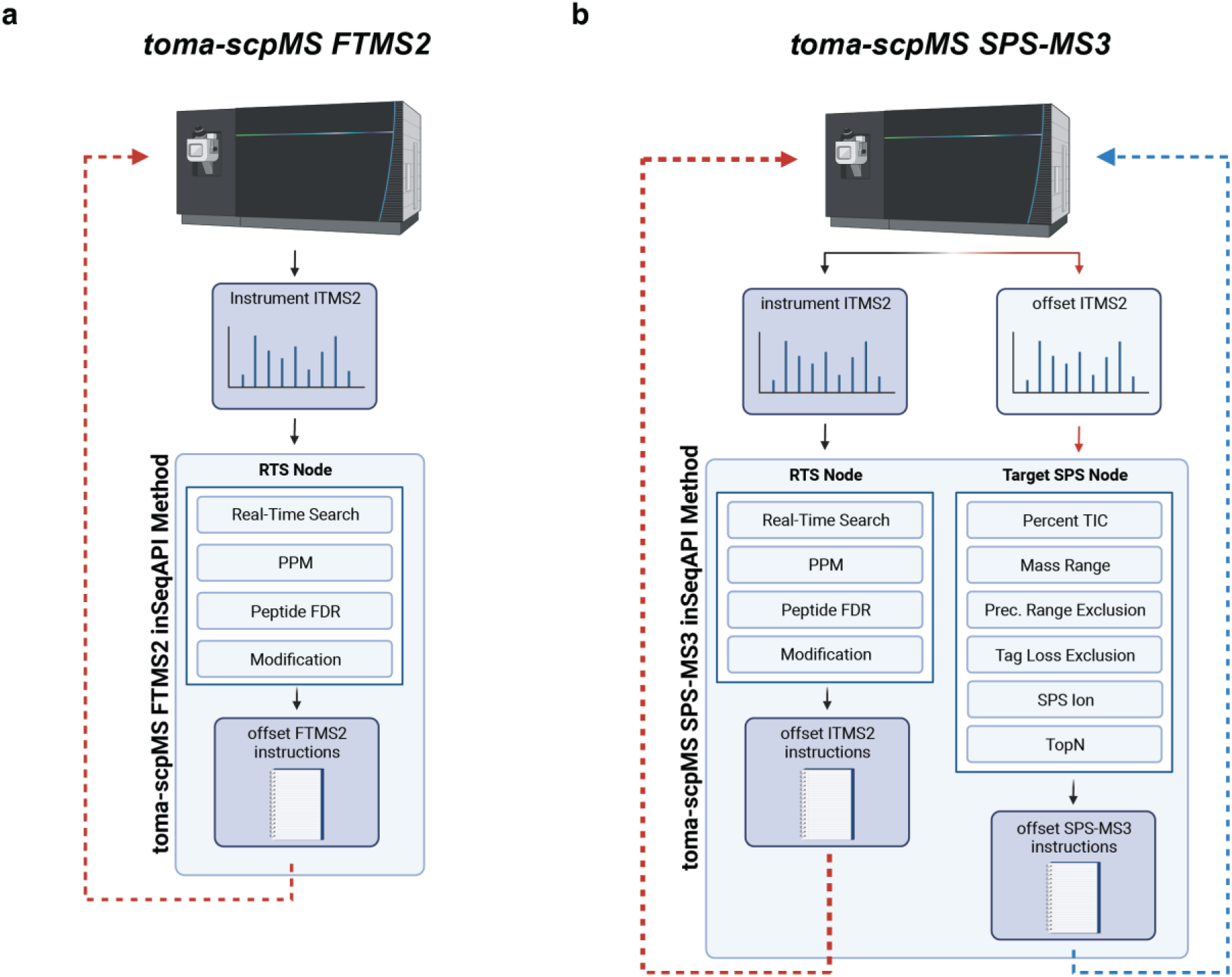
Detailed workflow for toma-scpMS methods within inSeqAPI. **a**, Workflow of toma-scpMS FTMS2 method within inSeqAPI. **b**, Workflow of toma-scpMS SPS-MS3 method within inSeqAPI.

**Supplementary Figure 2.**
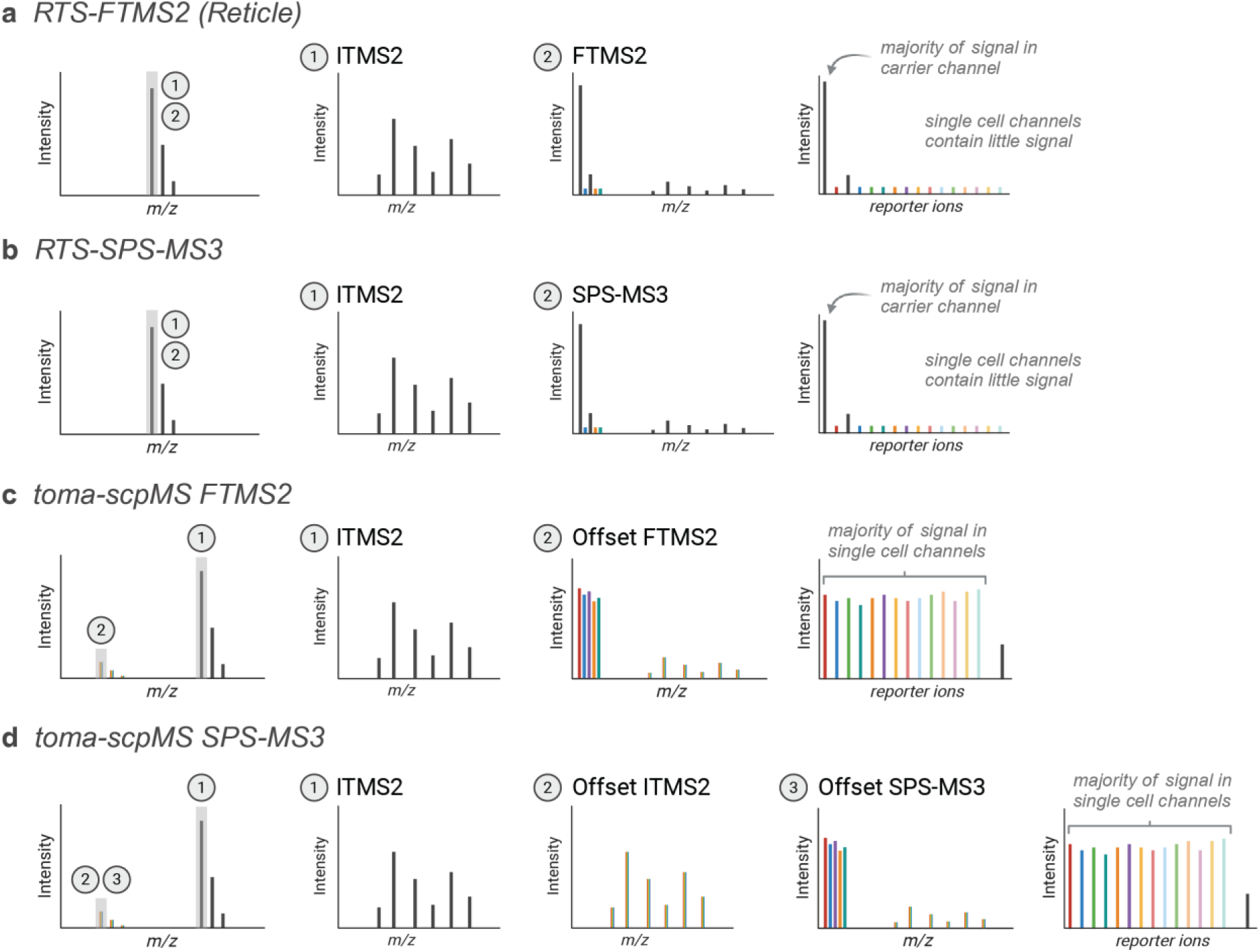
Method used for analysis of equimolar and mixed ratio samples. **a**, ‘RTS-FTMS2’, an ion trap scan followed by a real-time peptide identification triggered FTMS2. **b**, ‘RTS-SPS-MS3’ an ion trap scan followed by a real-time identification triggered SPS-MS3. **c**, ‘toma-scpMS FTMS2’, an ion trap scan of a carrier peptide followed by a real-time identification triggered FTMS2 on the single cell peptide. **d**, ‘toma-scpMS SPS-MS3’, an ion trap scan of a carrier peptide followed by a real-time identification triggered ion trap scan of a single cell peptide which was used to select SPS ions for a subsequent SPS-MS3 scan of the single cell peptide.

**Supplementary Figure 3.**
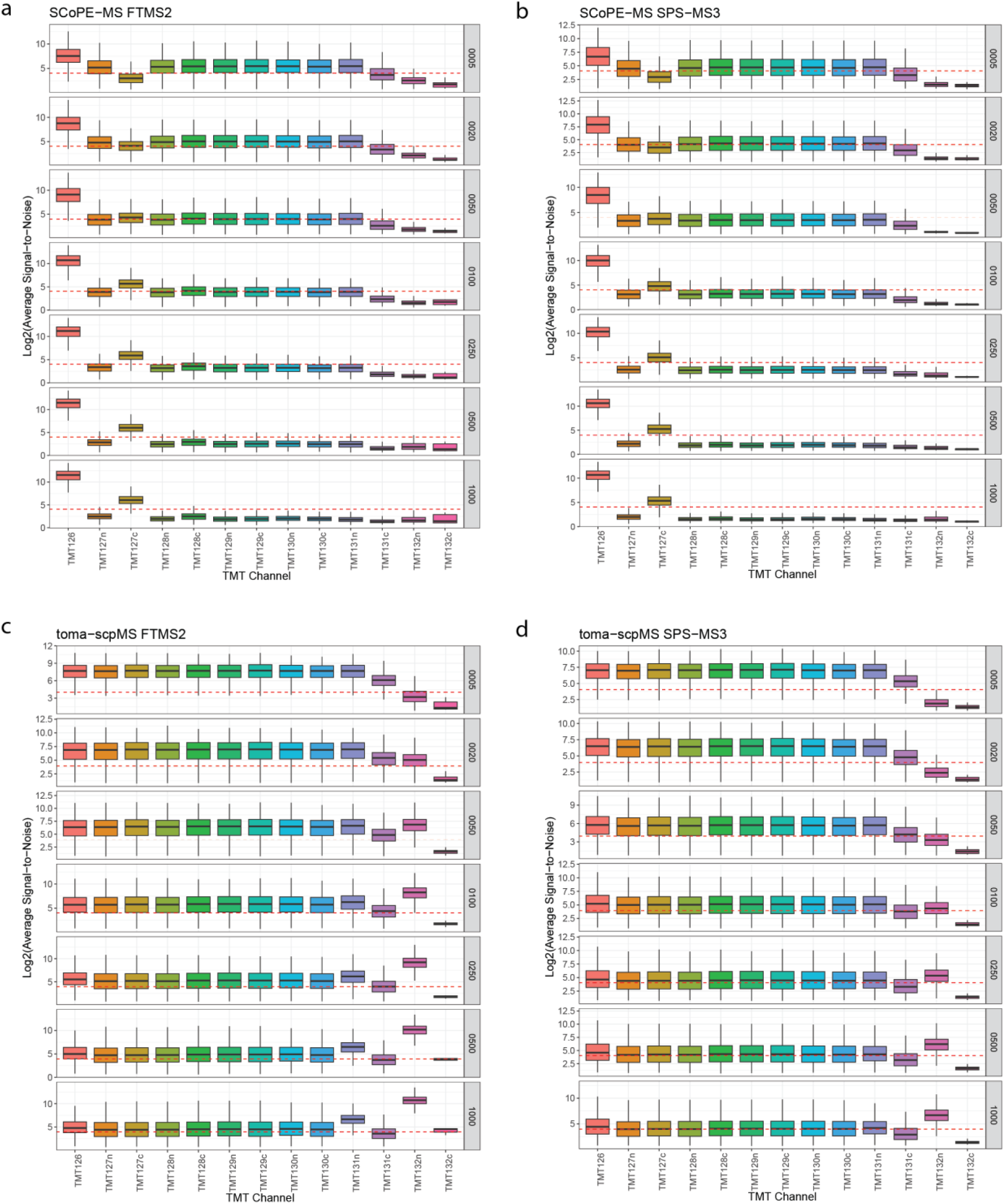
Signal-to-noise ratios for reporter ions within equimolar samples. Log2 transformation of the signal-to-noise of each reporter ion for SCoPE-MS FTMS2 quantification (**a**), SCoPE-MS SPS-MS3 quantification (**b**), toma-scpMS FTMS2 quantification (**c**), toma-scpMS SPS-MS3 quantification (**d**). The dotted line represents a QuantSN threshold of 16, the approximate threshold to be considered quantified.

**Supplementary Figure 4.**
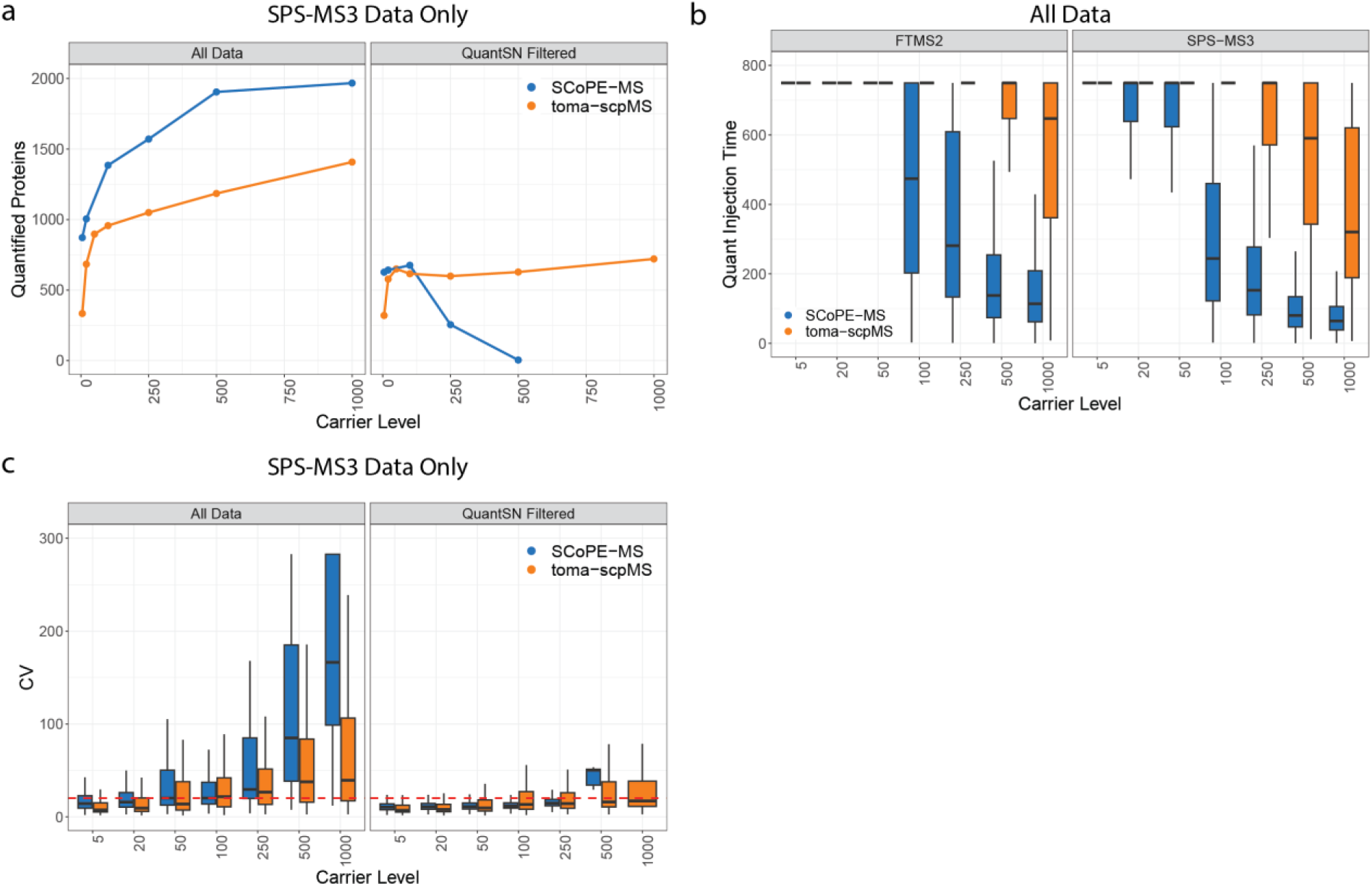
Supporting data for the analysis of equimolar mixtures with SCoPE-MS and toma-scpMS. **a**, The number of quantified proteins for toma-scpMS SPS-MS3 when all data are considered or only data that passes a QuantSN filter. **b,** Injection time in milliseconds (ms) for the quantification scan: FTMS2 for toma-scpMS FTMS2 and SPS-MS3 for toma-scpMS SPS-MS3. **c,** Coefficient of variation (CV) for toma-scpMS SPS-MS3 when all data is used (‘All Data’) or QuantSN filtered data are used (‘QuantSN Filtered’). For all box plots in c-d, the black line denotes the median while the box denotes the interquartile range and the whiskers the highest and lowest value, excluding outliers (not shown).

**Supplementary Figure 5.**
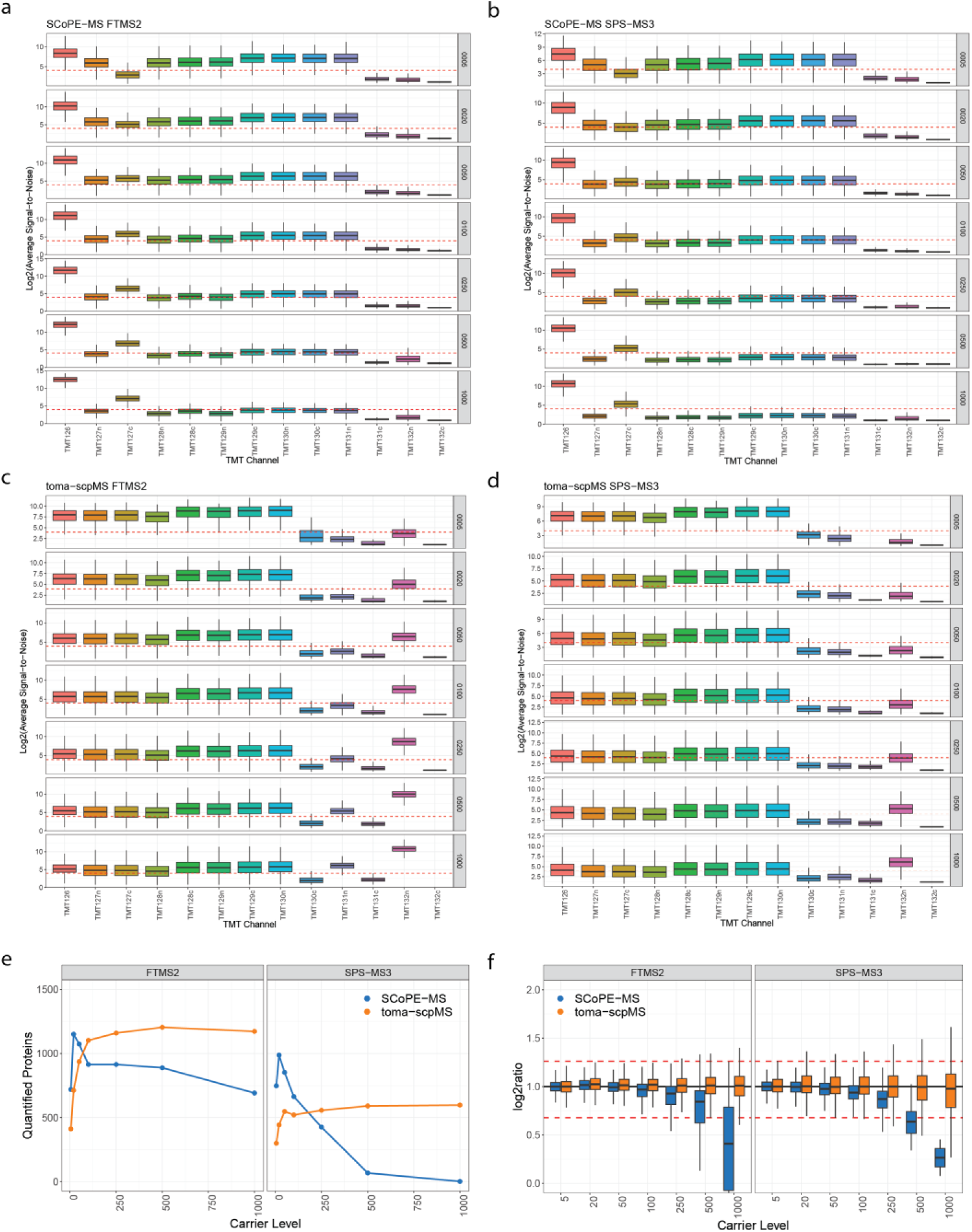
Supporting data for the analysis of mixed ratio samples by SCoPE-MS and toma-scpMS. Log2 transformation of the signal-to-noise of each reporter ion for SCoPE-MS FTMS2 quantification (**a**), SCoPE-MS SPS-MS3 quantification (**b**), toma-scpMS FTMS2 quantification (**c**), toma-scpMS SPS-MS3 quantification (**d**). **e**, The number of quantified proteins for the mixed ratio samples when the 128c channel is not considered for quantification. **f**, Distribution of measured protein ratios for the mixed ratio. The target ratio was 2 (log2ratio = 1, black horizontal line) and the red dotted lines represent a 20% measurement error. For all box plots, the black line denotes the median while the box denotes the interquartile range and the whiskers the highest and lowest value, excluding outliers (not shown). The dotted line represents a QuantSN threshold of 16, the approximate threshold to be considered quantified.

**Supplementary Figure 6.**
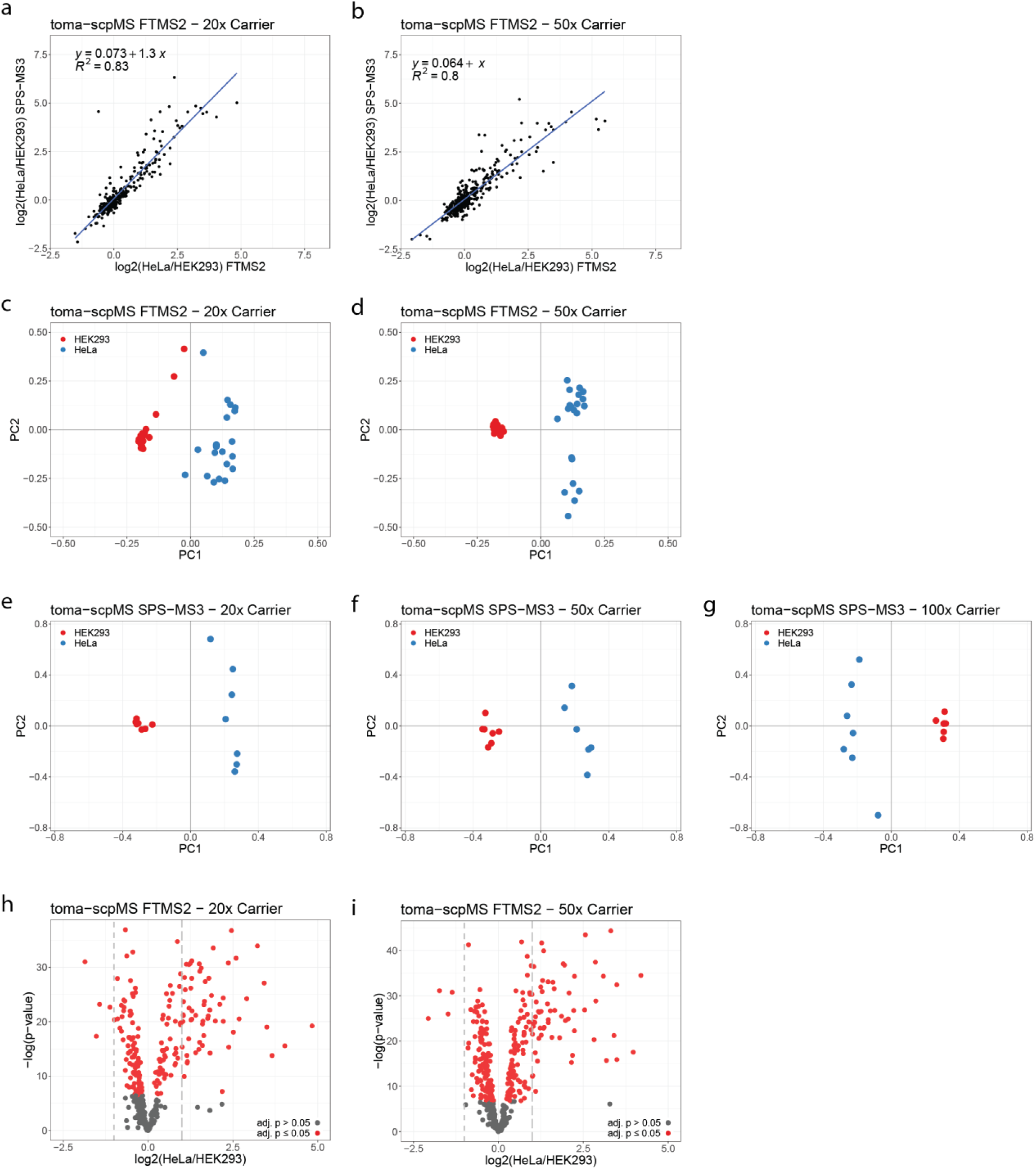
Supporting data for the analysis of single cells by toma-scpMS. **a-b**, Correlation of toma-scpMS FTMS2 and SPS-MS3 measurements for proteins quantified in the 20x and 50x carrier samplex using both methods. **c-g**, Principal component analysis based on proteins quantified across all three plexes differentiates cell types for noted acquisition method and carrier level. **h-i**, Volcano plot displaying the differential analysis of protein abundance across the two cell types for the noted acquisition methods and carrier level.

## Notes

https://massive.ucsd.edu/ProteoSAFe/dataset.jsp?accession=MSV000095629

